# Validated Determination of NRG1 Ig-like Domain Structure by Mass Spectrometry Coupled with Computational Modeling

**DOI:** 10.1101/2021.12.13.472484

**Authors:** Niloofar Abolhasani Khaje, Alexander Eletsky, Sarah E. Biehn, Charles K. Mobley, Monique J. Rogals, Yoonkyoo Kim, Sushil K. Mishra, Robert J. Doerksen, Steffen Lindert, James Prestegard, Joshua S. Sharp

**Affiliations:** Department of BioMolecular Sciences, University of Mississippi, University, MS 38677; Complex Carbohydrate Research Center, University of Georgia, Athens, GA 30602; Department of Chemistry and Biochemistry, Ohio State University, Columbus, OH 43210; Glycoscience Center of Research Excellence, University of Mississippi, University, MS 38677; Department of Chemistry and Biochemistry, University of Mississippi, University, MS 38677

## Abstract

High resolution hydroxyl radical protein footprinting (HR-HRPF) is a mass spectrometry-based method that measures the solvent exposure of multiple amino acids in a single experiment, offering constraints for experimentally-informed computational modeling. HR-HRPF-based modeling has previously been used to accurately model the structure of proteins of known structure, but the technique has never been used to determine the structure of a protein of unknown structure leaving questions of unintentional bias and applicability to unknown structures unresolved. Here, we present the use of HR-HRPF-based modeling to determine the structure of the Ig-like domain of NRG1, a protein with no close homolog of known structure. Independent determination of the protein structure by both HR-HRPF-based modeling and heteronuclear NMR was carried out, with results compared only after both processes were complete. The HR-HRPF-based model was highly similar to the lowest energy NMR model, with a backbone RMSD of 1.6 Å. To our knowledge, this is the first use of HR-HRPF-based modeling to determine a previously uncharacterized protein structure.

## Introduction

Mass spectrometry (MS) has rapidly gained in popularity not only in the identification and mass measurement of proteins, but in the characterization of protein higher order structure. Numerous MS-based technologies have been successfully used to characterize protein higher order structure, including hydrogen-deuterium exchange ^1^, limited proteolysis ^2^, chemical crosslinking ^3^and covalent labeling ^4^. Covalent labeling includes a number of techniques, all of which involve reaction of some reagent with amino acid side chains usually available on the surface of the folded protein. A variety of covalent labeling reagents have been used, including acylation reagents ^5^, diethylpyrocarbonate ^6^, carbenes ^7^, trifluoromethyl radicals ^8,9^ and iodine radicals ^10^. Here, we present an approach based on the use of hydroxyl radicals as a covalent labeling reagent. Hydroxyl radicals generate high quality data for a variety of amino acids, providing a generalizable probe for protein topography ^4,11–14^. We also demonstrate that this approach is capable of producing high quality reliable protein structures and validate this in a blind test against a parallel structure determination by NMR methods.

The approach we use begins with data from a technique known as hydroxyl radical protein footprinting (HRPF) ^15^. Hydroxyl radicals are useful and popular due to the wide variety of methods for *in situ* generation ^16–23^, broad reactivity ^13,14^, small size, hydrophilic nature, and well-characterized reaction pathways with various amino acids ^24^. Work from Chance and co-workers found that apparent rates of reaction could be correlated with average solvent accessible surface area (<SASA>) once the inherent rate of reaction of the amino acid was corrected using the free amino acids as a surrogate ^11,25^. Work from Sharp and co-workers confirmed these findings, further reporting that amino acids with lower inherent reactivity could display altered inherent reactivity based on sequence context ^12,26^. Sharp and co-workers further used amino acid-resolution HRPF (known as HR-HRPF) coupled with computational modeling to demonstrate the ability to differentiate between accurate computational models and inaccurate computational models, opening possibilities for using HR-HRPF data to determine protein structure^12^.

HR-HRPF data are then used to facilitate computational predictions of structure. The Lindert group developed the first software to use covalent labeling data in automated Rosetta protein structure prediction ^27,28^. Recently, Biehn and Lindert reported a more robust and computationally less expensive method for using HR-HRPF data to generate protein models using conical neighbor count instead of <SASA>, which successfully identified *ab initio* models of accurate atomic detail for three of the four benchmark proteins examined ^29^. However, while these studies indicate the potential of HR-HRPF for the determination of protein structure, no protein of unknown structure has had its structure determined solely using HR-HRPF data to inform computational modeling.

To accurately test the ability of HR-HRPF-based modeling to generate accurate structural models of novel proteins, we used the technology to determine the structure of the immunoglobulin-like domain (NRG1-Ig) of human neuregulin-1 (NRG1). NRG1 is a signaling glycoprotein that interacts with the ErbB/HER family of receptor tyrosine kinases via its EGF-like domain ^30–32^. NRG1-mediated signaling plays an important role in neuronal and cardiac development, and regulation of synaptic plasticity ^31–34^. Dysregulation of these signaling pathways is implicated in human disease, such as schizophrenia and certain forms of cancer ^35,36^. Due to a combination of alternative splicing and proteolytic processing, NRG1 exhibits a high diversity of isoforms, both soluble and membrane-bound, and a number of these isoforms include the Ig-like domain ^32,37^. In contrast to the EGF-like domain, the functional role of the NRG1-Ig domain is less well understood. It is believed to be involved in binding to heparan sulfate proteoglycans of the extracellular matrix ^38,39^, and there are reports that it can affect ErbB receptor activation ^40–42^.

In this manuscript, two teams worked independently to characterize the structure of NRG1-Ig. The first team used HR-HRPF to quantitatively measure topography of various amino acid side chains of the NRG1-Ig. Models of the protein were generated via Rosetta *ab initio* modeling, scored with the HRPF-guided Rosetta score term, then subjected to a Rosetta relaxation ensemble ^29^ from which a top scoring model was identified. Meanwhile, the second team determined the structure of NRG1 using standard heteronuclear solution NMR techniques. While all protein was expressed by the second team, no data were shared between groups to prevent any bias in structural characterization. After both teams had generated their structural models, the HR-HRPF constrained structure was compared to the NMR structure, to assess the accuracy of the HR-HRPF method. The results of this study serve as a rigorous and unbiased test of the ability of HR-HRPF to facilitate a reliable determination of soluble protein structures.

## Results and Discussion

### HR-HRPF of NRG1-Ig

Non-glycosylated versions of NRG1-Ig were expressed in *E. coli* (both with and without isotopic labeling) and purified as described in the Supplementary Information and **Figure S1**; structural homogeneity was verified by size exclusion chromatography and NMR. For MS studies, proteolytic digestion of NRG1-Ig was optimized for maximum sequence coverage after complete digestion to maximize HR-HRPF data and reproducibility. GluC was found to generate considerably higher sequence coverage than trypsin (**Figure S2**, Supplementary Information), with 98.3% coverage of the NRG1-Ig expressed sequence. GluC has also successfully been used in the past for HR-HRPF analysis, as the amino acids recognized by GluC are only minor oxidation targets. ^43^ Therefore, GluC was used for HR-HRPF analysis.

After purification and digestion optimization, multi-dose Fast Photochemical Oxidation of Proteins (FPOP) ^12,22,44^ was performed on NRG1-Ig material at natural isotopic abundance. A mixture of 10 μM NRG1-Ig, 17 mM glutamine, 1 mM adenine, 50 mM sodium phosphate, 2.2 mM Tris (pH 8.1), and hydrogen peroxide at 10 mM, 25 mM, 50 mM or 100 mM were used for FPOP labeling. Adenine dosimetry was measured for each experiment to determine delivered radical dose, in order to account for variability in radical generation or scavenging ^45^. A control for each FPOP peroxide concentration was conducted under the same conditions without laser irradiation to measure and correct for background oxidation.

Samples were then digested using our optimized GluC protocol. LC-MS/MS using electron transfer dissociation (ETD) was performed to measure the amount of oxidation at each amino acid for each oxidized peptide. Oxidation of twenty amino acids were measured (examples in **Figure 1**, with full data in **Figure S3**, Supplementary Information) and converted to the natural log of a protection factor (PF), defined as normalized relative intrinsic reactivity value for a particular free amino acid residue divided by the regression slope. 95% confidence intervals for regression slopes were used to represent uncertainty in PF measurement. Values measured for lnPF for all NRG1-Ig amino acids measured are given in **Figure S4** and **Table S1**, Supplementary Information.

**Figure 1.**
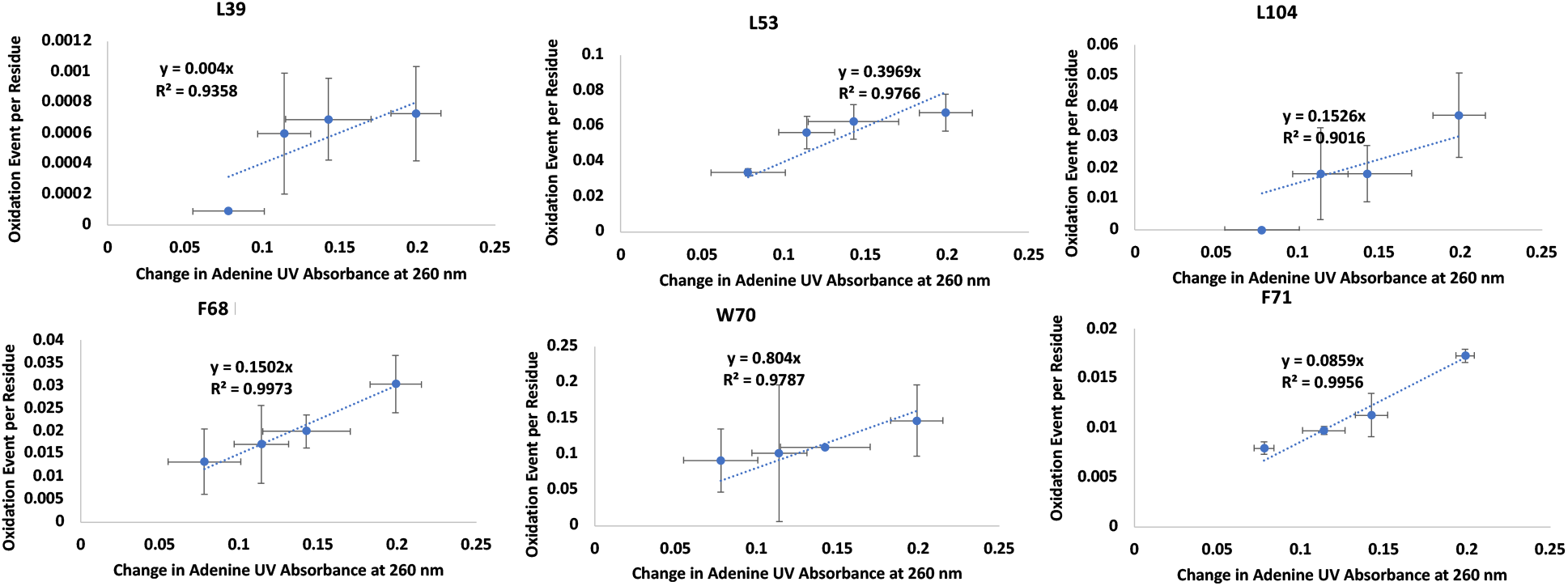
Measu red radical dose response rate of 6 amino acids used for structural modeling in NRG1-Ig via HR-HRPF. Each figure shows the calculated oxidation of each residue at 4 different hydrogen peroxide concentrations plotted against the change in adenine absorbance at 260 nm. The error bars represent one standard deviation from triplicate measurements for each data point. Each point represents the oxidation of one residue at a specific radical dose. The slopes of best-fit lines are radical dose responses.

### Solution NMR Structure of NRG1-Ig

Using a suite of standard multidimensional experiments (**Table S2**, Supplementary Information) we obtained nearly complete resonance assigments of ^1^H, ^13^C and ^15^N spins of the native polypeptide range (**Table S3, Fig. S5**, Supplementary Information). The only resonances we were unable to observe and assign were those of backbone ^1^H and ^15^N of Lys117. The ^13^C^α^ and ^13^C^β^ chemical shifts of the two cysteine residues were consistent with disulfide bond formation ^46^. Based on extensive chemical shift assignments and NOE data we obtained a well-defined solution NMR structure of NRG1-Ig (**Fig. S6, Table S3**, Supplementary Information). The fold of NRG1-Ig is typical of immunoglobulin-like domains, with a sandwich of two β-sheets stabilized by a disulfide bond. The smaller anti-parallel β-sheet consists of β-strands 41-58, 94-102, 86-91, while the second β-sheet consists of β-strands 77-72, 108-115, 120-130, 45-48 in a mixed topology with the last two stands running parallel. The only helical component is a single 310 turn at 104-106.

### Determining the Best Computational Models of NRG1-Ig

We employed our recent HRPF-guided Rosetta modeling protocol^29^ to predict the structure of NRG1-Ig. As per our published protocol, only lnPF values measured from Trp, Phe, Tyr, His and Leu were used. Our protocol used a HRPF score term, *hrf_dynamics*, that rewarded models demonstrating agreement with the FPOP labeling data. Additionally, we used Rosetta relaxation ensemble movers to sample protein flexibility. The output structures from the Rosetta mover protocol were referred to as mover models. Upon generation of 20,000 Rosetta *ab initio* models, we scored models with Rosetta’s score function (“Ref15”) (**Figure 2a**) and *hrf_dynamics* to determine a total score (**Figure 2b**). The 20 top scoring models were then used as inputs for the relaxation ensemble that generated thirty mover models per top scoring structure, leading to the addition of 600 models to be included in the model distribution (**Figure 2c**).

**Figure 2.**
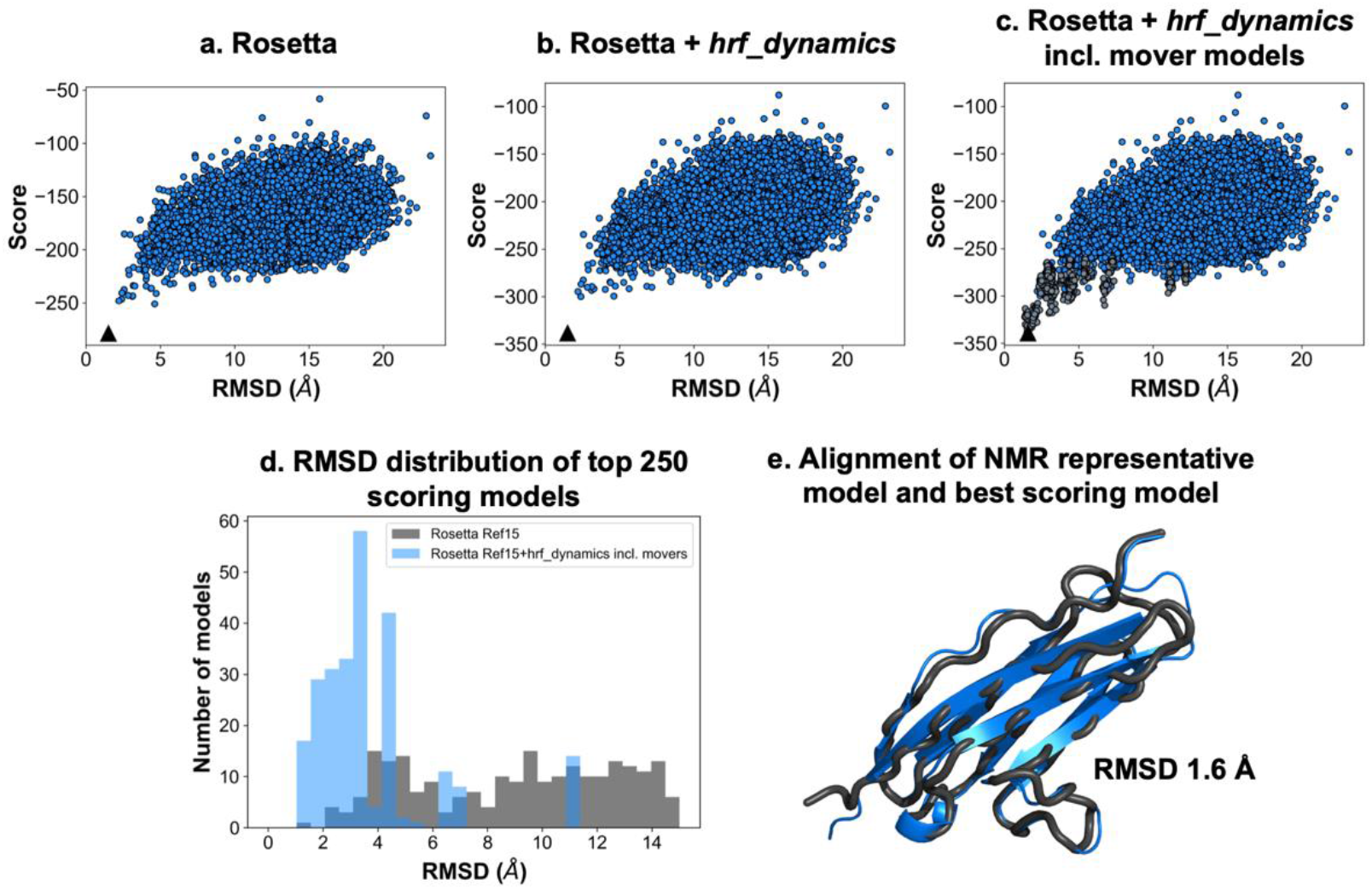
*Ab initio* modeling of NRG1-IG with relaxation ensemble and FPOP-guided scoring significantly enriched high-quality models. Score versus RMSD from NMR model 1 when **a**, scoring with Rosetta’s score function; **b**, scoring with Rosetta and *hrf_dynamics;* and **c**, scoring with Rosetta and *hrf_dynamics* including mover models (dark grey). Best scoring models are denoted by a black triangle. **d**, RMSD histograms for top 250 scoring models when scoring with Rosetta (grey) versus Rosetta and *hrf_dynamics* including mover models (blue). Bin widths were maintained at 0.5 Å. **e**, Alignment of NMR model 1 (black) with the top scoring model identified from our HRPF-guided and mover model protocol (blue). The RMSD to the NMR model was determined to be 1.6 Å.

Upon examination of the 250 top scoring models when scoring with Rosetta versus scoring with Rosetta and *hrf_dynamics* including mover models, we observed a decrease in the average root mean squared deviation (RMSD) and an increase in the percentage of models with RMSDs under 5 Å (**Figure 2d**). The average RMSD of the top 250 models when scoring with Rosetta was 9.5 Å, which improved to 3.8 Å when scoring with FPOP data and including mover models. When scoring with Rosetta, 21% of the top 250 models had RMSDs below 5 Å. This improved with *hrf_dynamics* usage and mover model generation to 94% of models having RMSDs under 5 Å. When scoring both mover models and *ab initio* structures with our score term, we identified one of the generated mover models as the best scoring model. Our best scoring model exhibited backbone RMSD of 1.6 Å to the determined NMR structure of NRG1-Ig (**Figure 2e**).

The use of optimized conical neighbor count (the number of neighbors within the vicinity of a residue based on distance and angle contributions) ^29^ to apply FPOP data to computational models is a significant alternative to the use of amino acid solvent accessible surface area data. The correlation between the HR-HRPF lnPF results for NRG1-Ig optimized conical neighbor count from the lowest energy NMR structure was consistent with correlations previously reported for model protein structures ^11,12,47^. The subset of amino acids considered here are robust regardless of the method of hydroxyl radical generation or amino acid-level quantitation, and no bias was introduced due to over-fitting to known structures (**Figure 3**). Overall, employment of the relaxation ensemble to generate mover models resulted in a significant enrichment of accurate, high-quality, low-RMSD models in this blind prediction effort. We concluded that usage of our FPOP-guided and relaxation ensemble method can increase confidence in model selection for other structure prediction efforts.

**Figure 3.**
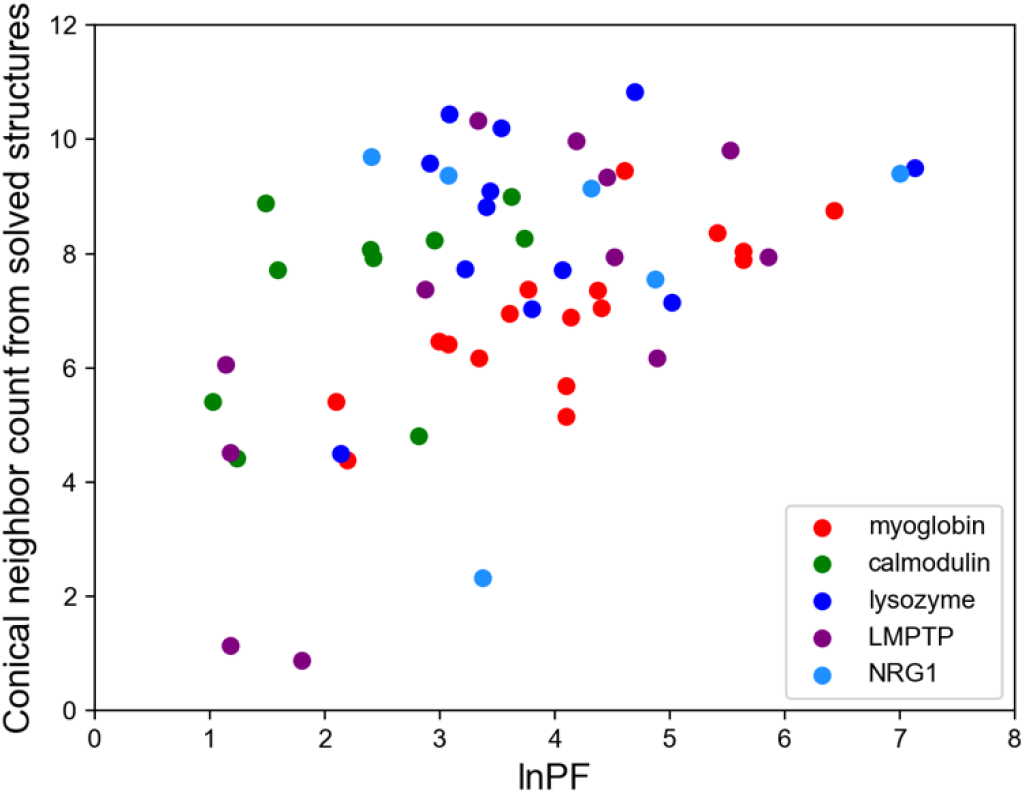
Correlation between HR-HRPF lnPF and conical neighbor count. The correlation measured for NRG1 performed blinded to the NMR structure (cyan) was consistent with those reported for proteins of known structure.

## Conclusion

In this work, we tested the ability of HR-HRPF combined with conical neighbor count computational modeling to generate accurate, reliable structural models of a protein of truly unknown structure, NRG1-Ig. We were able to greatly increase the reliability of Rosetta modeling by application of HR-HRPF data, generating a final model with a backbone RMSD of <2 Å from the lowest energy NMR model, and with a large increase in model reliability. As the NRG1-Ig structure was unknown when HR-HRPF was performed and the NMR structure was determined independent of the HR-HRPF group, we have excluded any possibility ofconfirmation bias in our experimental design. The consistency of our results with previous work published on proteins of known structure shown in **Figure 3** reveals a lack of confirmation bias in these previous results, and indicates no clear difference in accuracy based on the method of radical generation or amino acid-level oxidation quantification for the subset of amino acids used here (Trp, Phe, Tyr, His and Leu).

Our results as independently confirmed in a blind study by established NMR techniques demonstrate that HR-HRPF combined with conical neighbor count computational modeling is not just a tool for examining relative changes in protein topography, but is a structural biology tool that generates experimentally informed computational models of protein structure that are accurate and reliable. With the rise in computational tools for structural prediction, including the recently released AlphaFold ^48,49^, there is a need for flexible experimental methods to validate predicted structures. HR-HRPF has no theoretical limitations on the size or dynamics of measured protein structures, is compatible with protein complexes and can be carried out using microgram quantities of protein. Given the flexibility and low sample requirements of HR-HRPF compared with traditional high-resolution structural biology techniques, this methodology can play a significant role in the validation of computational structures, as well as in the generation of accurate and reliable structural models when computational methods fail. The application of HR-HRPF constraints could also support modern computational methods for predicting structures of protein complexes^50^, which currently have suspected issues regarding comprehensiveness and accuracy in certain cases that could be greatly remedied through the application of experimental HR-HRPF results. Future work examining the ability of HR-HRPF combined with conical neighbor count to correctly identify domain-domain contacts and orientation will be important for developing the application of HR-HRPF combined with conical neighbor count to address challenging problems in multi-domain protein structural biology.

## Materials and Methods

### Expression & purification of NRG1-Ig

A pET-21b(+) plasmid containing a TEV-cleavable N-terminal His-tag and a 100 residue fragment comprising the NRG1-Ig domain (residues 34-133 of the UniProt Q02297 sequence) was purchased from GenScript (US distribution, Piscataway, NJ). This plasmid was transformed into BL21(DE3) *E. coli* cells (New England Biolabs, Ipswich, MA) using standard protocols. Transformed cells were applied onto LB agar plate with ampicillin followed by overnight incubation at 37 °C. A single colony was used to inoculate a 10 mL LB media with carbenicillin and incubated overnight at 37 °C. Cells were pelleted at 2000 g and resuspended in 3 ml of M9 media. Resuspended cells (600 ul) were used to inoculate a 50 ml M9 culture and incubated at 37 C until OD600=0.8.

Glycerol stocks of transformed cells were then used to inoculate 10 mL of LB starter culture, followed by overnight incubation at 35 °C. Cells were then pelleted and resuspended in 1 L of LB media, and incubated again at 35 °C. To produce NRG1-Ig at natural isotopic abundance, the 1 L culture was induced with 1mM IPTG after reaching OD_600_ of ~0.6; 3h after induction cells were harvested by centrifugation at 2,500 x g and frozen. For stable isotope labeled samples, the 1L LB culture was instead pelleted upon reaching OD_600_ of ~0.8, and the cell pellet was resuspended in 0.5L of M9 minimal media containing ^15^NH_2_Cl with either ^13^C-glucose or 5% ^13^C-glucose (Cambridge Isotope Laboratories). Incubation continued for about 1hr at 35 C when expression was induced with 1 mM IPTG. Cells were harvested after ~3hrs and frozen.

In both cases, thawed cells were resuspended in lysis buffer (20mM Tris pH 8.1, 300mM NaCl, and 1mM TCEP with protease inhibitors) at 4°C and lysed using a French-press. The resulting lysate was centrifuged at 12,000 x g, and the pellet fraction containing inclusion bodies was resuspended in denaturing buffer (6M Urea, 300mM NaCl, 1mM TCEP, 6mM imidazole and 20mM Tris pH 8.1) at 4 °C using either handheld or electric tissue homogenizer. NRG1-Ig was purified under denaturing conditions via immobilized metal affinity chromatography (IMAC) using NGC system (Bio-Rad) equipped with a 10 mL Co-NTA column. Elution of NRG-Ig1 was accomplished with a linear gradient beginning with 3% Buffer A (6M Urea, 20mM Tris pH 8.1 at 4 °C, 300mM NaCl, 1mM TCEP) and ending with 100% Buffer B (6M Urea, 20mM Tris, pH 8.1 at 4 °C, 200mM imidazole, and 300mM NaCl). The recovered U-^15^N,^13^C, U-^15^N, 5%-^13^C, and natural abundance NRG1-Ig fractions were sealed in in dialysis tubing (Spectrapor, 6-8 kDa) and refolded by dialysis at 4 °C in four steps against a refolding buffer (20mM Tris pH 8.1 at 4 °C, 300mM NaCl). The refolding buffer was supplemented with 0.1 mM TCEP and 50uM ethylenediaminetetraacetic acid (EDTA) for the first dialysis stem. U-^15^N,5% U-^15^N,^13^C- and U-^15^N,5%-^13^C-labeled NRG1-Ig were subsequently exchanged into NMR buffer (20 mM sodium phosphate pH 6.5, 100mM NaCl) using 0.5 ml Amicon microconcentrators. U-^15^N,^13^C NMR samples (~35 μl in 1.7 capillary NMR tube), NRG1-Ig NC(I) and NRG1-Ig NC(II), consisted of 0.45 mM and 2.0 mM NRG1-Ig, respectively, with 0.05% sodium aside, 4 μM sodium trimethylsilylpropanesulfonate (DSS) and 7% D_2_O. U-^15^N,5% ^13^C NMR sample, NRG1-Ig NC5 (~40 μl in 1.7 capillary NMR tube), was prepared in the original TRIS refolding buffer, with 0.05% sodium aside, 5 μM sodium trimethylsilylpropanesulfonate (DSS) and 7% D_2_O. For HRPF studies, the NRG1-Ig batch without isotope labeling was subjected to an additional purification stem on a Waters BEH SEC Column, 125 Å, 1.7 μm, 4.6 mm * 300 mm using a Thermo Fisher Dionex 3000 HPLC system. The running buffer was 20 mM Tris at pH 8.1 with 300 mM NaCl using an isocratic gradient.

### Multi-Dose FPOP and NRG1-Ig Digestion

FPOP was performed in triplicate for neuregulin using a 248 nm COMPex Pro 102 high pulse energy excimer KrF laser in the presence of various hydrogen peroxide concentrations (10 mM, 25 mM, 50 mM, and 100 mM) ^12^ The experiment was done in triplicate for each hydrogen peroxide concentration. For FPOP, samples were prepared by mixing NRG1-Ig to the final concentration of 10μM in 50mM sodium phosphate (Sigma-Aldrich), 17 mM glutamine (Acros Organics) and 1 mM adenine (Acros Organics) as a radical dosimeter ^45^. Freshly prepared hydrogen peroxide (Fisher Scientific) at 4 different concentrations (10 mM, 25 mM, 50 mM, and 100 mM) was added to each sample prior laser exposure. A total volume of 20 μl of sample flowed through the excitation capillary at 17.34 ul/min. The nominal laser fluence at the plane of the excitation capillary was at 9.82 mJ/mm^2^ with 15% exclusion volume. After laser irradiation, the samples were quenched in 25 ul quenching buffer containing 50 nM catalase (Sigma-Aldrich) and 20 mM methionine amide (Bachem). A control sample for each hydrogen peroxide concentration was done in triplicate with the laser turned off. After laser exposure, we measured the changes in adenine UV absorbance of each oxidized sample as compared to each control at 265 nm using a nanodrop spectrophotometer. This represents the effective radical dose delivered to the protein^12^.

After quenching, the oxidized and control samples were denatured and reduced at 95 °C for 30 minutes in the presence of 5.5 mM DTT (Soltec Ventures). After denaturation, the samples were put on ice for 2 minutes. More sodium phosphate buffer at pH 6 was added to keep its concentration at 30 mM prior to GluC addition. GluC (Promega Corp) was added in 1:20 enzyme: protein mass ratio. The samples were digested overnight for 14 hours.

### C18 RPLC-MSMS C18

LC-MS/MS was done using an Acclaim PepMap 100 C18 nanocolumn (0.075 mm × 150 mm, 2 μm particle size, 100 Å pore size, Thermo Fisher Scientific) coupled to a 300 μm i.d. ×5 mm C18 PepMap 100 trap column with 5 μm particle size (Thermo Fisher Scientific) to desalt and concentrate the samples before loading onto the C18 nanocolumn for separation. LC-MS buffers were made using all LC/MS grade solvents (Fisher Scientific). The capillary pump was used to load the samples onto the C18 trap column using buffer C (water + 0.05% TFA) and buffer D (acetonitrile + 0.05% TFA). We used a nanopump for chromatographic separation using mobile phase E (water+ 0.1% formic acid) and mobile phase F (acetonitrile+ 0.1% formic acid). Initially, the samples were loaded onto the C18 trap column in 98% C, 2% D at 5 μl/min for 6 minutes. The trap column was then switched inline with the nanocolumn and trapped peptides were back-eluted onto the nanocolumn using the nanopump. Elution started by increasing buffer F in a linear gradient from 2% to 40% (with the balance as buffer E) over 22 minutes. The gradient was then ramped up to 95% F over 5 minutes and held isocratic for 3 minutes to wash the column. Buffer F was then decreased to 2% over 1 minute and held isocratic for 6 minutes to re-equilibrate the column for the next run. The samples were eluted directly into a nanospray source of a Thermo Fusion Tribrid orbitrap, where the spray voltage was set at 2600 V and ion transfer tube temperature at 300 °C. A full MS scan was obtained from 150 to 2000 m/z. CID and ETD was performed every 2 seconds on precursor ions of +2 charge and greater for peptide identification and sequence coverage analysis. For ions with +2 charge state, ETD was performed with 20% EThcD SA collision energy to increase ETD fragmentation. The orbitrap resolution for both ETD and EThcD was 30000 with Automatic Gain Control target at 5e4 and maximum injection time of 100 ms.

### Peptide and Amino Acid level Oxidation Analysis

Byonic version v2.10.5 (Protein Metrics) was used to identify NRG1-Ig peptide sequences using the NRG1-Ig protein sequence described in Figure S2 (Supplementary Information). For all peptides detected, the major oxidation products detected were net additions of one or more oxygen atoms. In order to calculate average oxidation events per peptide, the area under the curve for peaks of unoxidized and oxidized peptides was used according to **Equation 1**. Briefly, the oxidation events per peptide were calculated by summing the intensity (*I*) of each peptide oxidation product multiplied by the number of oxidation events on the peptide required to generate that product and divided by the sum of *I* for all oxidized and unoxidized versions of that peptide, as shown in **Equation 1**. *P* represents the average oxidation events per peptide, and *I* is the area under the curve for peaks of oxidized and unoxidized peptides.

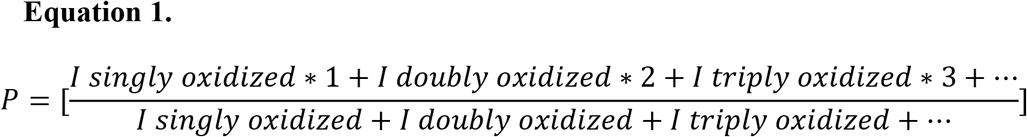

The amount of oxidation at residue level quantitation in a peptide was determined by the fragment ion (z or c ion) intensities of the peptide ETD fragmentation. The oxidation fraction of a given z or c ion was calculated by dividing the oxidized sequence ion intensity by the sum of the intensity of the corresponding oxidized and unoxidized sequence ion in a particular oxidized peptide. The relative oxidation fraction of each product ion *f*(Z_i_) was calculated using **Equation 3.2** where I(Z_i_) is the intensity of the designated production summed across all spectra.

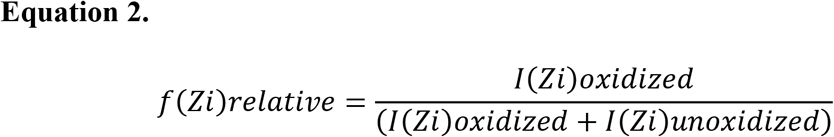

The absolute amount of oxidation of a given amino acid was determined by multiplying the average oxidation event of a peptide by the absolute fractional oxidation of the corresponding sequence ions. As shown in **Equation 3**, *P* is the average oxidation event per peptide calculated from **Equation 1**, and the term in brackets is the fractional difference of two adjacent sequence ions, f(Z_i_) and f(Z_i–1_). In cases where ETD fragmentation ions are not adjacent in sequence, fractional oxidation for multiple contiguous residues within the peptide was calculated by using non-adjacent ETD fragments in **Equation 3**.

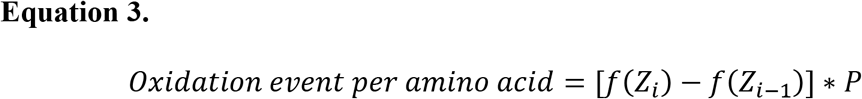

In order to take background oxidation into account, the oxidation event of each residue was calculated by subtracting the oxidation event of the same residue under control conditions from its oxidation event in the oxidized sample.

Natural Log Protection Factor (ln(PF)) was calculated using **Equation 4** where *R_i_*, represents the amino acid intrinsic reactivity for residue *i* while *Slope*_i_ represents the experimentally determined radical dose response for residue *i*.

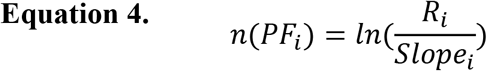

### Structural Modeling

Using Rosetta’s *AbInitioRelax* protocol, the sequence of NRG1-Ig (adapted from UniProt Q02297), and fragment libraries obtained from the Robetta server, 20,000 *ab initio* models of NRG1-Ig were built.^51–55^ No FPOP data were included during model generation. Models were scored with the “Ref15” Rosetta score function. Per-residue FPOP data were converted into the natural log of the protection factor (lnPF).^25,28,29^ The lnPF values were supplied to the *hrf_dynamics* term, and models were scored based on their agreement with the labeling data.^29^ The summed per-residue *hrf_dynamics* score used a weight of 9.0, as described previously. The total score was determined by adding the Rosetta and *hrf_dynamics* scores. Models were ranked by total score. The twenty top-scoring models were then used as input for mover model generation with the Rosetta relaxation ensemble, as described previously.^29^ For each of the top-scoring structures, thirty mover models were obtained. The six hundred mover models were scored with Rosetta and *hrf_dynamics* and then included in the *ab initio* model distribution. The best scoring model was selected as our blind prediction for the NRG1-Ig structure, for comparison with the NMR structure. For comparison between HR-HRPF and NMR structural models, Cα RMSD values with no outlier rejection were calculated with Rosetta.

### NRG1-Ig NMR Spectroscopy and Structure Calculation

NMR data collection, processing, resonance assignment, and structure calculation followed the protocols of Northeast Structural Genomics Consortium (NESG Wiki, http://www.nmr2.buffalo.edu/nesg.wiki/Main_Page). NMR spectra (**Table S2**, Supplementary Information) for NRG1-Ig samples were acquired at 25 °C on an AVANCE NEO 800 MHz spectrometer (Bruker BioSpin) equipped with a 1.7 mm TCI ^1^H(^13^C,^15^N) cryogenic probe. All spectra were Fourier-transformed using Topspin v4 (Bruker Biospin), except non-uniformly sampled 3D HBHA(CO)NH, which was reconstructed using Smile ^56^ and Fourier-transformed with NMRPipe ^57^. ^1^H chemical shifts were referenced relative to 4,4-dimethyl-4-silapentane-1-sulfonic acid (DSS), and ^13^C and ^15^N chemical shifts were referenced indirectly via gyromagnetic ratios. Visualization and analysis of NMR spectra, NOE peak picking and integration were performed with the program CARA ^58^. Automated assignment of backbone ^1^H, ^15^N, ^13^CO, ^13^C^α^ and ^13^C^β^ resonances was obtained with AutoAssign ^59^ followed by interactive validation and completion. Side-chain resonances were assigned interactively using 3D (H)CCH and 3D ^13^C/^15^N-edited [^1^H,^1^H] NOESY spectra. Stereospecific assignments of Leu and Val isopropyl groups were obtained based on positive versus negative peak intensities in the 2D [^13^C,^1^H] constant-time HSQC (CT-HSQC) acquired for NRG1-Ig NC5, as described previously ^60^. Stereospecific assignment of Asn and Gln CONH_2_ groups were determined from relative NOE peak intensities.

Structure calculation and automatic NOE peak assignment was performed iteratively using CYANA v 3.98.13 ^61,62^ and ASDP v1.0 ^63^. Constraints for backbone φ, ψ and side-chain χ_1_ dihedral angles were derived using TALOS-N ^64^, and those that were consistent with the initial structural models were used in subsequent structure calculation steps. NOE peaks with matching unambiguous assignments from CYANA and ASDP were manually checked and refined for consistency with NOE spectra and distance constraint violations, and then used to optimize NOE distance calibration function. Assignments of these peaks were kept fixed during final structure calculation with CYANA. Stereospecific assignment of CH2 groups was performed iteratively using the GLOMSA module of CYANA. Of 100 calculated conformers, 20 conformers with the lowest target function values were further refined in explicit water bath using CNS ^65^ as previously described ^66^ with distance constraints relaxed by 5%. The quality of NRG1-Ig structure models was analyzed with PSVS ^67^, and the resulting statistics are summarized in **Table S3**, Supplementary Information. Software used for NMR data analysis and structure calculation was accessed via NMRBox ^68^. Atomic coordinates, structural restraints, assigned NMR chemical shifts and NOE peaklists were deposited in the Protein Data Bank (PDB ID 7SJL) and BioMagResBank (accession code 30960).

## Acknowledgements

This work was supported by the National Institute of General Medical Sciences (R01GM127267). S.M. and R.J.D. acknowledge support from the Glycoscience Center of Research Excellence (NIH P20GM130460) and J.H.P. acknowledges support from (R01GM033225). This study made use of NMRbox: National Center for Biomolecular NMR Data Processing and Analysis, a Biomedical Technology Research Resource (BTRR), which is supported by NIH grant P41GM111135 (NIGMS). Modeling work in this publication was supported by NSF (CHE 1750666) to S.L.

