## Supplementary Information for "Validated Determination of NRG1 Ig-like Domain Structure by Mass Spectrometry Coupled with Computational Modeling"

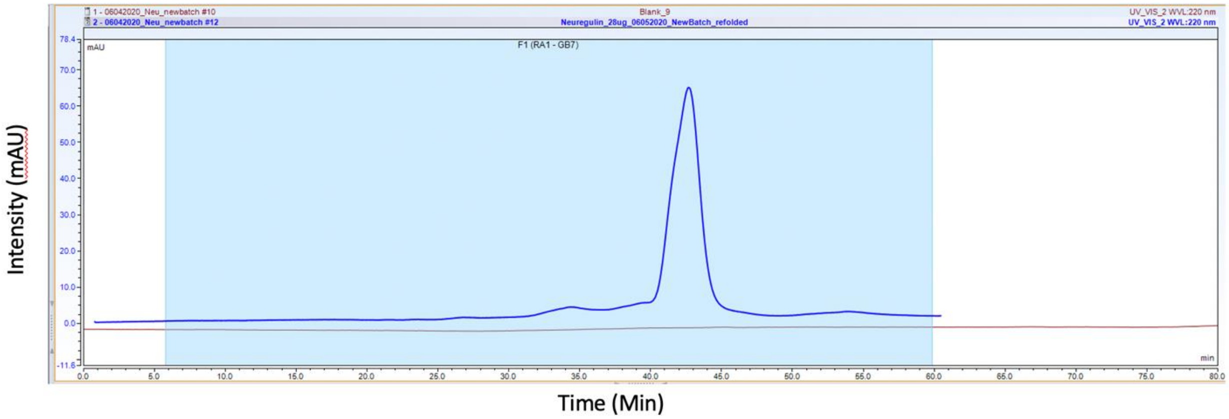


**Figure S1. NRG1-Ig Analysis by SEC.** Brown line represents the absorbance at 220 nm ($A_{220}$) of the blank run before running neuregulin. Blue line represents the $A_{220}$ of NRG1-Ig, indicating the presence of one main conformation within the protein sample. Bottom-up MS/MS analysis of the major peak (retention time = 42-43 min) verified the identity as NRG1-Ig (data not shown).

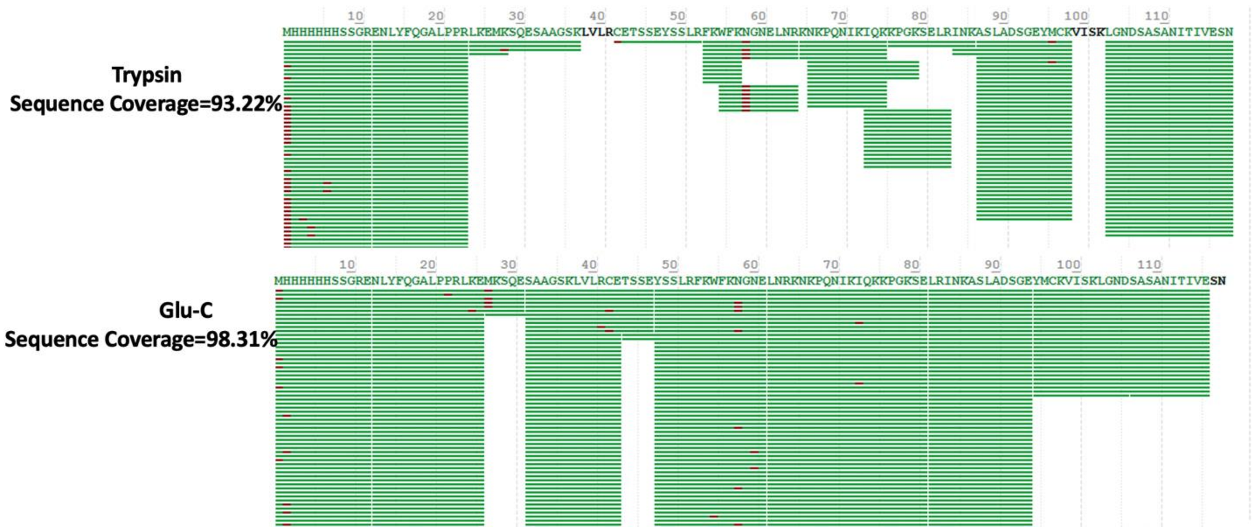


**Figure S2. Sequence Coverage of Neuregulin using Trypsin and GluC enzymes.** (A) Sequence coverage of neuregulin digested by trypsin. (B) Sequence coverage of neuregulin digested by GluC. Each horizontal bar represents an MS/MS spectrum assigned to the peptide covered by the width of the bar. Red spots represent chemical modifications to the peptide assigned by Byonic.

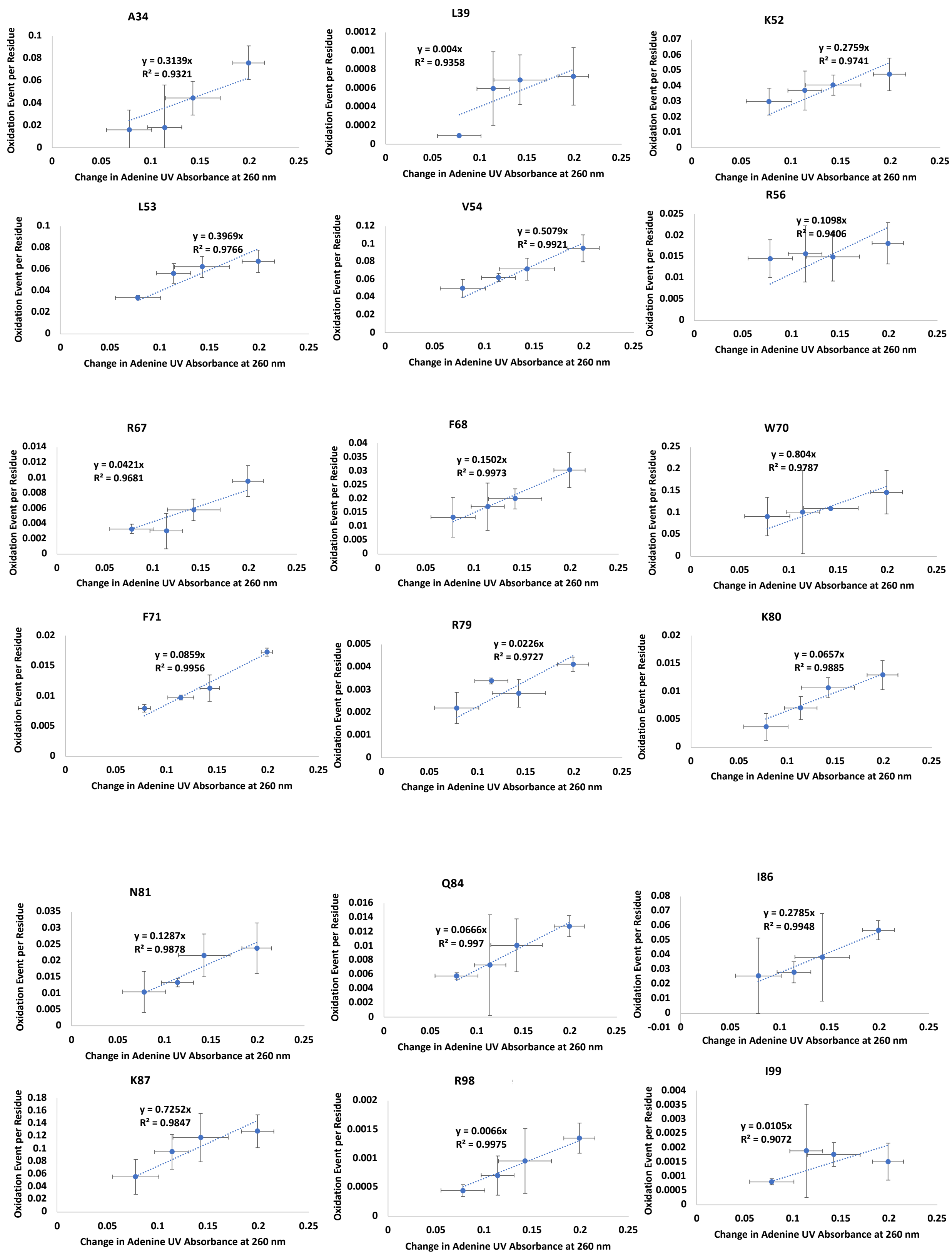
A34
y = 0.3139x
R² = 0.9321
L39
y = 0.004x
R² = 0.9358
K52
y = 0.2759x
R² = 0.9741
L53
y = 0.3969x
R² = 0.9766
V54
y = 0.5079x
R² = 0.9921
R56
y = 0.1098x
R² = 0.9406
R67
y = 0.0421x
R² = 0.9681
F68
y = 0.1502x
R² = 0.9973
W70
y = 0.804x
R² = 0.9787
F71
y = 0.0859x
R² = 0.9956
R79
y = 0.0226x
R² = 0.9727
K80
y = 0.0657x
R² = 0.9885
N81
y = 0.1287x
R² = 0.9878
Q84
y = 0.0666x
R² = 0.997
I86
y = 0.2785x
R² = 0.9948
K87
y = 0.7252x
R² = 0.9847
R98
y = 0.0066x
R² = 0.9975
I99
y = 0.0105x
R² = 0.9072
Oxidation Event per Residue
Change in Adenine UV Absorbance at 260 nm

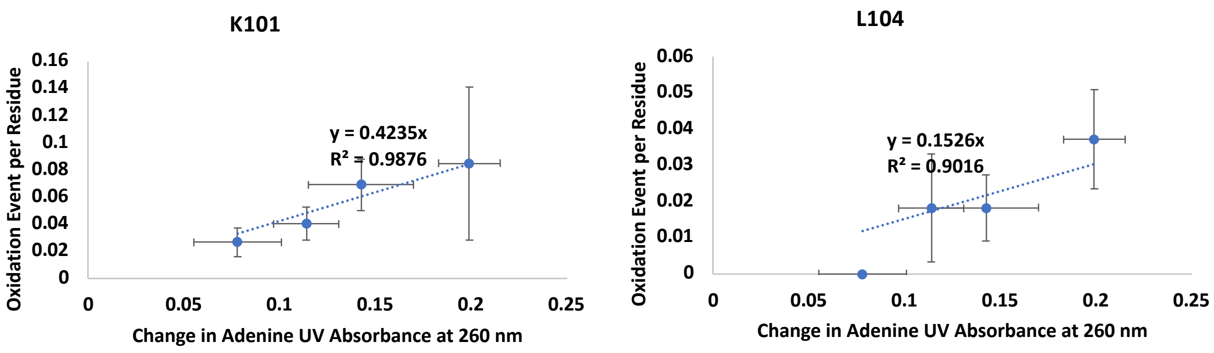


**Figure S3. Measured radical dose response rate of all NRG1 amino acids in native form via HR-HRPF.** Each figure is the calculated oxidation event of each residue at 4 different hydrogen peroxide concentrations plotted against the changes in adenine absorbance at 260 nm. The error bars represent one standard deviation from triplicate measurements for each data point. Each point represents the oxidation event of one each residue at a specific radical dose. The slope of this correlation is radical dose response.

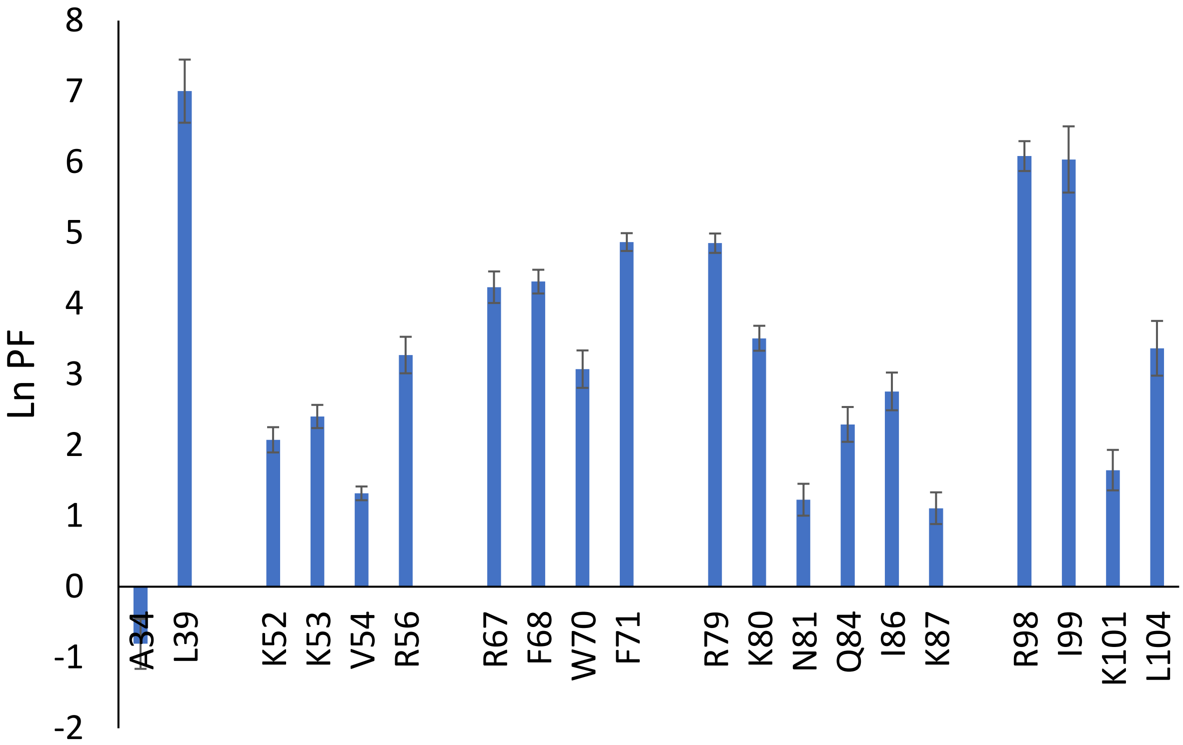


**Figure S4. lnPF for each amino acid measured in NRG1.** Error bars represent the 95% confidence interval based on linear regression analysis performed in **Figure 1** and **Figure S3**.

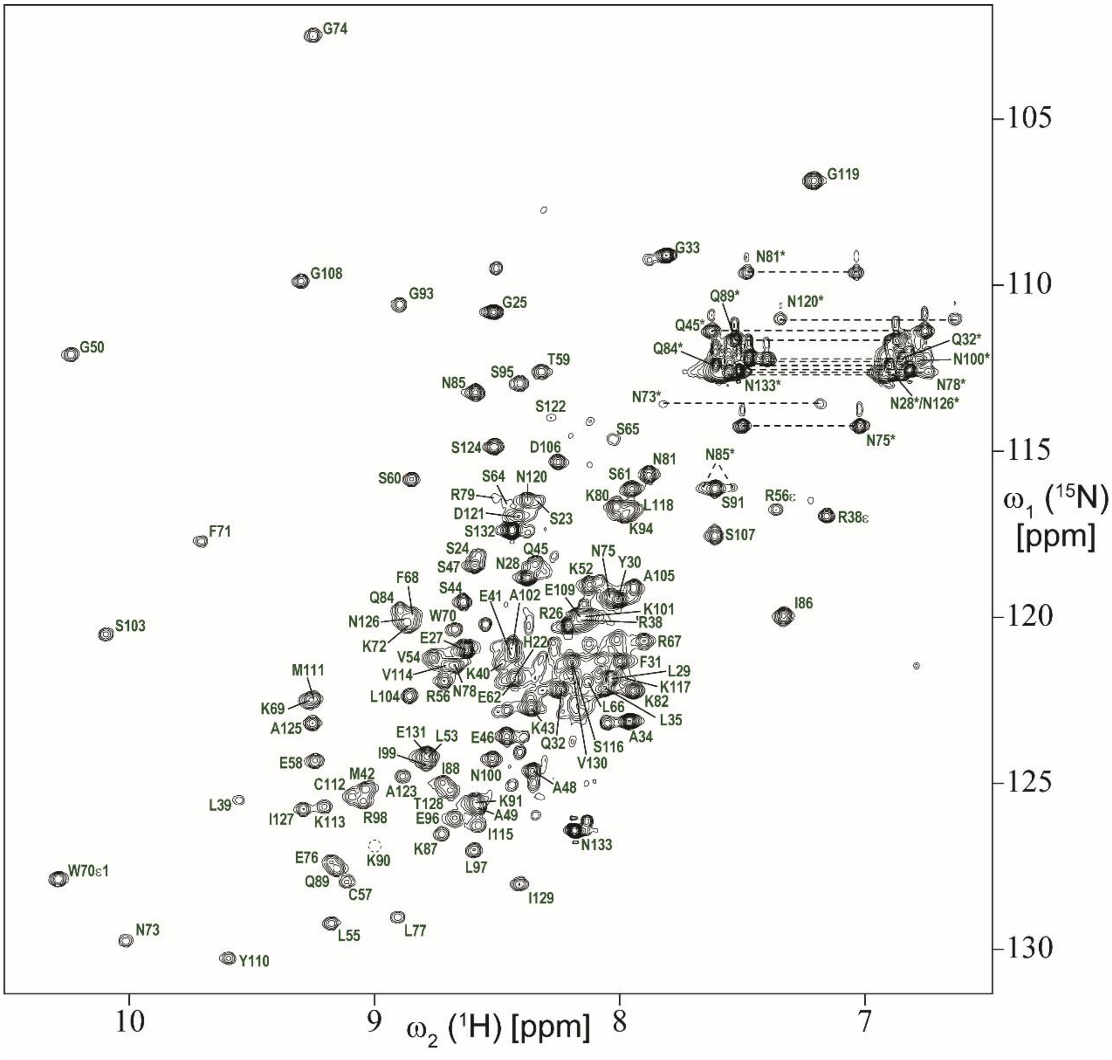


**Figure S5.** 2D [$^{15}$N, $^{1}$H] HSQC spectrum of NRG1-Ig NC(II) sample acquired at 800 MHz magnetic field strength. Peak assignments are labeled in green with residue numbering corresponding to UniProt Q02297. Side chain resonances of Asn and Gln resonances are indicated with asterisks, also shown are side-chain resonances of Trp70 Hε1/Nε1, as well as Hε/Nε of Arg38 and Arg56. The latter two signals are 'aliased' into the observable region from their true $^{15}$N chemical shift positions. Backbone amide signal of Lys90 is below the lowest displayed contour level, and its position is indicated by a dotted circle.

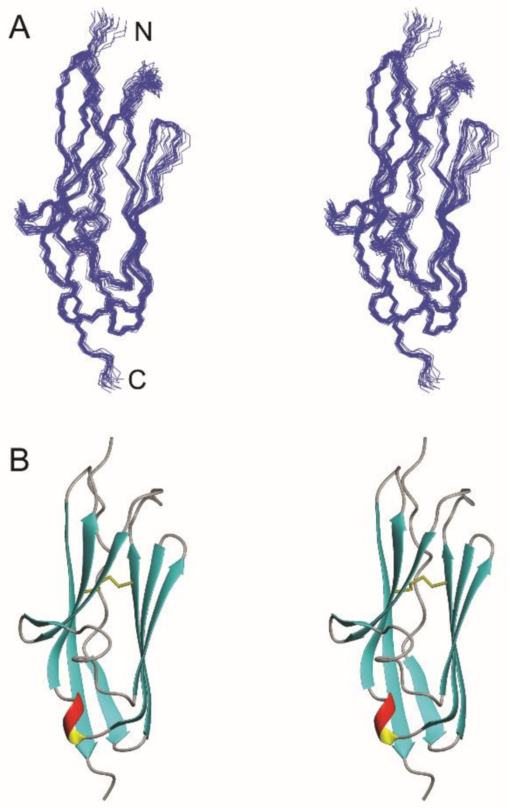


**Figure S6.** Stereo pair representation of solution NMR structure of NRG1-Ig. Only the native residues 34-133 are shown. **A.** Line plot of the 20 representative conformers superimposed for minimal RMSD of the $C^{\alpha}$ atoms of ordered residues. N- and C-termini are indicated. **B.** Ribbon diagram of the lowest-energy conformer showing the regular secondary structure elements. The disulfide bond linking the two β-sheets is shown as yellow sticks.

**Table S1. lnPF for each amino acid measured in NRG1.**

| Amino Acid | lnPF ± 95% CI |
|---|---|
| A34 | -0.81 ± 0.35 |
| L39* | 7.00 ± 0.45 |
| K52 | 2.08 ± 0.18 |
| L53* | 2.41 ± 0.16 |
| V54 | 1.32 ± 0.10 |
| R56 | 3.27 ± 0.26 |
| R67 | 4.23 ± 0.22 |
| F68* | 4.31 ± 0.17 |
| W70* | 3.07 ± 0.26 |
| F71* | 4.87 ± 0.13 |
| R79 | 4.85 ± 0.14 |
| K80 | 3.51 ± 0.18 |
| N81 | 1.23 ± 0.23 |
| Q84 | 2.29 ± 0.25 |
| I86 | 2.76 ± 0.27 |
| K87 | 1.11 ± 0.22 |
| R98 | 6.09 ± 0.21 |
| I99 | 6.04 ± 0.47 |
| K101 | 1.65 ± 0.29 |
| L104* | 3.37 ± 0.39 |

***Amino acids used for *hrpf_dynamics* scoring**

**Table S2. List of multidimensional NMR experiments used for resonance assignment and structure determination**

| Sample | Experiment |
|---|---|
| NRG1-Ig NC 2mM | 2D [$^{15}$N, $^{1}$H] HSQC |
| | 2D [$^{13}$C, $^{1}$H] CT-HSQC (aliphatic) |
| | 2D [$^{13}$C, $^{1}$H] CT-HSQC (aromatic) |
| | 3D (HACA)CONH |
| | 3D HNCACB |
| | 3D HBHA(CO)NH (NUS) |
| | 3D (H)CCH-COSY aromatic |
| | 3D (H)CCH-COSY aliphatic |
| | 3D (H)CCH-TOCSY aliphatic, 16 ms mixing time |
| | 3D $^{13}$C/$^{15}$N-edited [$^{1}$H,$^{1}$H] NOESY, 80 ms mixing time |
| NRG1-Ig NC 450uM | 3D HNCO |
| | 3D CBCA(CO)NH |
| NRG1-Ig NC5 | 2D [$^{13}$C, $^{1}$H] CT-HSQC methyl, 28ms CT delay |

**Table S3. NRG1-Ig structure statistics**

| | |
|---|---|
| **Resonance assignment completeness[a)] [%]** | |
| Backbone | 99.6 |
| Side-chain | 100.0 |
| **Conformation-restricting distance constraints[b)]** | |
| Intra-residue [i = j] | 367 |
| Sequential [\|i – j\| = 1] | 606 |
| Medium-range [1 < \|i – j < 5] | 309 |
| Long-range [\|i – j] ≥ 5 | 1150 |
| Total | 2432 |
| **Dihedral angle constraints ($\varphi/\psi/\chi_1$)** | 90/90/29 |
| **NOE constraints per restrained residue** | 24.0 |
| Of those, long range | 10.5 |
| **Average number of dihedral angle constraint violations per conformer > 10°** | 0 |
| **Average RMSD from mean coordinates[c)] [Å]** | |
| Backbone atoms | 0.5 |
| Heavy atoms | 0.9 |
| **Global quality scores[c)] (raw / Z-score)** | |
| PROCHECK G-factor (phi-psi) | -0.59/-2.01 |
| PROCHECK G-factor (all) | -0.37/-2.19 |
| Molprobity clash score | 5.64/0.56 |
| ProsaII | 0.25/-1.65 |
| Verify3D | 0.16/-4.82 |
| **Molprobity[3] Ramachandran summary[c)] [%]** | |
| Most favored regions | 96.4 |
| Additionally allowed regions | 3.3 |
| Disallowed regions | 0.4 |
| **CYANA target function [Å$^2$] (average over 20 conformers)** | 0.26 |
| **RPF analysis** | |
| Recall/Precision/F-measure | 0.959/0.918/0.938 |
| DP-score | 0.846 |

a) Commonly observed protein NMR resonances. Excludes residues of the N-terminal purification tag, as well as side-chain amino groups of Lys, side-chain guanidinium groups of Arg, carboxyl groups of Asp and Glu, and hydroxyl $^1$H of Ser, Thr and Tyr, and non-protonated aromatic $^{13}$C.
b) Calculated with PSVS v1.5
c) Ordered residue ranges: 34-61,66-132.
